# Preprinting Microbiology

**DOI:** 10.1101/110858

**Authors:** Patrick D. Schloss

## Abstract

The field of microbiology has experienced significant growth due to transformative advances in technology and the influx of scientists driven by a curiosity to understand how microbes sustain myriad biochemical processes that maintain the Earth. With this explosion in scientific output, a significant bottleneck has been the ability to rapidly disseminate new knowledge to peers and the public. Preprints have emerged as a tool that a growing number of microbiologists are using to overcome this bottleneck. Posting preprints can help to transparently recruit a more diverse pool of reviewers prior to submitting to a journal for formal peer-review. Although use of preprints is still limited in the biological sciences, early indications are that preprints are a robust tool that can complement and enhance peer-reviewed publications. As publishing moves to embrace advances in internet technology, there are many opportunities for preprints and peer-reviewed journals to coexist in the same ecosystem.

## Background

A preprint is an interim research product that is made publicly available before going through an official peer-review process with the goals of soliciting feedback, accelerating dissemination of results, establishing priority, and publicizing negative results (1–5). Authors can post their manuscript to a preprint server for others to read, share, and comment. In the 1960s, Information Exchange Groups were the first formal attempt to broadly disseminate paper-based preprints among physicists and biologists (6, 7). Although the biological community’s commitment to preprints waned by 1967, the physics community persisted and eventually adopted what is now the *arXiv* (pronounced “archive”) preprint server that was hosted at the Los Alamos National Laboratories from 1991 to 1999 and then at Cornell University (8). For some physicists and mathematicians, posting a preprint to *arXiv* optionally followed by submission to a peer-reviewed journal has become a standard publication pathway. Although *arXiv* has hosted a number of computational biology preprints, the server has not drawn widespread attention from biologists. Among proponents of *arXiv*, preprints have aided in the development of research communication by accelerating the release of science and helping authors reach a wider audience for critique and establishment of priority (9). Considering the broadening adoption of preprints among microbiologists, I sought to explore the specific uses of and concerns regarding preprints.

## Landscape of preprint servers

In 2013, two preprint servers, the *bioRxiv* (pronounced “bio-archive”) and *PeerJ Preprints*, were launched as preprint servers for biologists that would parallel *arXiv* (10). According to information provided on the *bioRxiv* and *PeerJ Preprints* websites and my personal experiences, both platforms offer similar features: preprint posting is free; each preprint receives a digital object identifier (DOI) that facilitates the ability to cite preprints in other scholarly work; if the preprint is ever published, the preprint is linked to the published version; the submission process for both options is relatively simple allowing authors to upload a PDF version of their preprint and supplemental materials; preprints are typically publicly available in about 24 hours; they have built-in venues for authors to discuss their research with people who leave comments on the manuscript; preprints undergo a basic screening process to remove submissions with offensive or non-scientific content; and the sites provide article-level metrics indicating the number of times an abstract has been accessed or the preprint has been downloaded. There are several important differences between the two options. First, *PeerJ Prints* is a for-profit organization and *bioRxiv* is a non-profit organization sponsored by Cold Spring Harbor Laboratory. This difference can be meaningful to authors since some journals, including the American Society for Microbiology (ASM) Journals, will only accept submissions that have been posted on preprint servers hosted by not-for-profit organizations (11). Second, preprints at *PeerJ Preprints* are posted under the Creative Commons Attribution License (CC-BY) and *bioRxiv* preprints can be posted under one of four CC-BY licenses or with no permission for reuse. This can be relevant for authors hoping to submit their work to *Proceedings of the National Academy of Sciences* as the journal will not consider manuscripts posted as preprints under a CC-BY license. The NIH encourages authors to post preprints under the CC-BY or public domain licenses (5). The flexibility of the *bioRxiv* licensing empowers authors to choose the model that best suits them, while ensuring the rapid posting of their research results; however, it is important to provide clear information to authors on the legal and practical tradeoffs of each option. A cosmetic, but still relevant difference is the layout and feel of the two websites. Compared to the *bioRxiv* site (Figure S1), the *PeerJ Preprint* site is more fluid, gives readers the ability to “follow” a preprint, and provides better access to article keywords and the ability to search preprints (Figure S2). With broader acceptance of preprints by traditional journals, many journals, including all of the ASM journals, have established mechanisms to directly submit manuscripts that are posted as preprints on *bioRxiv*. The only direct submission mechanism for manuscripts submitted at *PeerJ Preprint* is to the *PeerJ* journal. In many ways, preprint servers have taken on the feel of a journal. As adoption of this approach expands, it is likely that the features of these sites will continue to improve. It is also worth noting that numerous other opportunities exist for other forms of interim research products (e.g. blog posts, videos, protocols, etc.) to obtain DOIs that make the work citable. As these possibilities increase, the preprint landscape risks becoming fractured.

One solution to the fracturing of the preprint landscape would be the creation of indexing sites that allow a user to easily search for content across multiple preprint servers. Several examples of these efforts already exist and it is likely that these interfaces and their ability to span the landscape will improve. For example, although Google Scholar includes preprints hosted at *bioRxiv* and *PeerJ Preprints* in their search results, PubMed and Web of Science do not. A relatively new example of what this might look like is PrePubMed (12), which seeks to index preprints from numerous sources.

A more organized effort is being initiated with funding through ASAPbio to create a “Central Service” that would aggregate preprints in the life sciences (13). As preprint servers and other content providers begin to look and act like traditional journals by incorporating features and interfaces, it is important to value the strength of the preprint - that of an interim research product that is nimble and quickly posted. It is therefore essential to balance the requirements placed on authors for features associated with preprints with the efficiency of the preprint format.

## Specific challenges for microbiology

Although preprints offer an efficient and novel venue for disseminating microbiology research, there are several considerations that the scientific community and those that oversee preprint servers must consider. It is critical that assurances be given that policies are in place to address these issues and that these policies are made transparent. First, attention has to be given to dual use research of concern (DURC) since microbiology-related research could offer insights to individuals seeking to engage in inappropriate activities. Second, for researchers engaging in research that involves human subjects and other vertebrates, it is critical that assurances be made that institutional oversight committees have been consulted and have approved of the research. Third, there is significant concern regarding researchers disclosing potential conflicts of interest that could affect a project’s experimental design, analysis, and interpretation of results. Finally, recent expansions in scientific publishing have revealed numerous cases of plagiarism or misconduct. Again, while hoping to maintain the efficiency of the preprint format, traditional microbiology journals have screening procedures and oversight committees that address these issues. Similar efforts need to be implemented by preprint servers. As preprint usage continues to expand many of these problems may also grow similar to the experiences within the traditional publishing industry has expanded.

## Acceptance of preprints by journals

An early controversy encountered by researchers interested in posting their work as preprints as a stage in disseminating their research was whether it constituted prior publication (14). The broad consensus of the International Committee of Medical Journal Editors and numerous journals is that preprints do not constitute prior publication (15). This consensus is reflected in the current policies of journals that commonly publish microbiology research including those published by ASM, the Microbiology Society, International Society for Microbial Ecology, PLOS, the *Proceedings of the National Academy of Science*, *Science*, *Nature*, *Journal of Infectious Diseases*, and Cell press. Each take a generally permissive stance towards posting of preprints prior to submission. Comprehensive lists of journals’ attitudes towards preprints are available online and are regularly updated (16, 17). Considering the relatively fluid nature of many of these policies and the journals’ specific policies, prospective authors should be aware of the positions taken by the journals where they may eventually submit their work.

## Preprints and peer-review

The use of preprints for citations in other scientific reports and grant proposals has recently been called into question (18). It is important to note that the peer-review process was adapted to the technologies and trends that have evolved over the past 100 years. The formal peer-review system that most journals currently use was not developed until the end of the 1800s with the advent of typewriters and carbon paper (19). Editorial decisions were typically made by a single person or a committee (i.e. the editorial board) who had an expertise that covered the scope of the journal. As science became more specialized, new journals would form to support and provide a source of validation to the new specialty. The growth in science in the mid 1900s resulted in a shift from journals struggling to find sufficient numbers of manuscript to publish to having too many manuscripts submitted. It has been argued that the widespread adoption of decentralized peer-review was due to the increased specialization and to deal with the large number of manuscript submissions (20). Peer-review did not achieve widespread use at many journals, including the *Journal of Bacteriology*, until the 1940s and 1950s. Thus the “tradition” of peer-review is only 70 years old. Given the rapid advances in communication technology and even greater specialization within microbiology, it is worth pondering whether the current scientific publishing system and peer-review system, in particular, need to continue to adapt with our science.

Communicating research has traditionally been done within research group meetings, departmental seminars, conferences, and as publications. Along this continuum, there is an assumption that the quality of the science has been improved because it has been vetted by more experts in the field. The public dissemination of one’s research is a critical component of the scientific method. By describing their research, scientists subject their work to formal and informal peer-review. Their research is scrutinized, praised, and probed to identify questions that help seed the next iteration of the scientific method. A common critique of more modern approaches to publishing has been an inability to assess the quality of the science without the validation of peer-review. Attached to assertions of the validity of the research has been assertions of the impact and robustness of the research. These are all quality assessments that many acknowledge are difficult to assess by the traditional peer-review process. This has led to some journals, most notably *PLOS ONE*, calling for referees to place a reduced emphasis on the perceived impact or significance of the work. It has also led to the call for replacing or complementing pre-publication peer-review with post-publication peer-review using PubMed Commons, PubPeer, journal-based discussion forums, F1000Research, and other mechanisms. Alas if scientists are going to depend on post-publication peer-review or informal methods of peer-review for documents like preprints, they must be willing to provide constructive feedback on the work of others.

## Preprints have the potential to change the advancement of science

Preprints are often viewed as existing in a state of scientific limbo. As noted above, they represent a formal communication, but an interim one, not officially published. As the use of preprints grows and scientists’ perceptions of preprints matures, there are a number of issues that will need to be addressed.

First, a common concern is that if a researcher posts their work as a preprint, it will be “scooped” by another researcher and the preprint author will lose their ability to claim primacy or their ability to publish the work in a journal. Considering the preprint is a citable work with a DOI, it would, in fact, be the preprint author that scooped the second. Furthermore, a preprint could prevent getting scooped since a preprint would indicate to others in the field that the work had already been done, which would prevent wasted time and effort. The use of preprints uncouples the communication of the discovery from the relevance of the discovery, which will come later based on peer-review, comments from other scientists at meetings or online, and eventually citations. A growing number of scientific societies and journals, including ASM view preprints as citable and as having a legitimate claim to primacy (1, 21–23); however, it remains to be determined whether the journals will stand by these policies. Some scientists worry that with such protection a researcher can make a claim without valid data to support their claims (3). This is possible; however, it is also the responsibility of the scientific community to utilize the peer-review mechanisms that are available to comment on those preprints pointing out methodological problems or to indicate that they are speaking beyond the data. As preprints gain broader adoption, the tension between establishing primacy and the completeness of the preprint may test the policies of preprint-friendly journals.

A second area of concern is whether a preprint can be used to support a grant proposal. Given the length limitations placed on grant proposals by funding agencies, there is a push to cite previous work to indicate a research team’s competence in an area or to provide preliminary data. The National Institutes of Health (NIH) recently released a notice clarifying their position on the use of preprints and synthesizing feedback that they received as part of a request for information (5). In this notice, the NIH indicated that preprints can be cited anywhere that other research is cited including research plans, bibliographies, biosketches, and progress reports. Some fear that the use of preprints will allow scientists to circumvent page limits by posting preliminary manuscripts (24). One would hope that both consumers of preprints and grant proposal reviewers would be able to differentiate between someone trying to game the system and someone that is using preprints as a mechanism to improve their science. This would be greatly facilitated by following the NIH recommendation of using preprints as evidence for research progress, but providing an indication that the preprints are not peer-reviewed publications (5). This would help review panels in rendering their decisions and help authors substantiate their preliminary data.

A third concern is what role preprints should have in assessing a scientist’s productivity. Clearly use of publication metrics as an indicator of a scientist’s productivity and impact is a contentious topic without even discussing the role of preprints. Regardless, given the propensity for researchers to list manuscripts as being “in preparation” or “in review” on an application or curriculum vitae, listing them instead as preprints that can be reviewed by a committee would significantly enhance an application and a reviewer’s ability to judge the application. In fact, several funding agencies including the NIH, Wellcome Trust, UK Medical Research Council encouraging fellowship applicants to include preprints in their materials (5).

Beyond these concerns, preprints are also causing some to change their publication goals. Some authors are explicitly stating that a preprint will not be submitted to a journal (25). Although these authors may be a minority of those who post preprints, such an approach may be attractive to those who need to cite a report of a brief research communication, a critique of another publication, or negative results. It is clear that the adoption of preprints will challenge how scientists interact and evaluate each other’s work. There is great potential to empower researchers by controlling when a citable piece of work is made public.

## Microbiology anecdotes

The peer-review editorial process can be lengthy and adversarial. Because preprints are public and freely available they represent a rapid and potentially collaborative method for disseminating research. Several anecdotes from the microbiology literature are emblematic of benefits of the rapid release cycle that is inherent in the use of preprints.

First, preprints have proven useful for rapidly disseminating results for disease outbreaks and new technologies. Prior to the recent Zika virus outbreak there were approximately 50 peer-reviewed publications that touched on the biology and epidemiology of the virus; as of April 2017 the number of Zika virus-related peer-reviewed publications was over 2,300. During the recent outbreak, more than 150 Zika virus-related preprints have been posted at *bioRxiv*. Any manuscript that was formally published went through several month delays in releasing information to health care workers, the public, and scientists needing to learn new methods to study a previously obscure virus. In contrast, those that posted their work as a preprint were able to disseminate their methods and results instantly. Another interesting use of preprints to disseminate new information about Zika virus has been the posting of a preprint describing the Zika virus outbreak in the US Virgin Islands that will be continually updated as new data and analyses are performed (26). Over the last several years there have also been rapid advances in DNA sequencing technologies that have fundamentally changed how microbial science is performed. One notable technology, the MinIon sequencing platform from Oxford Nanopore, has received considerable attention from researchers who have posted more than 110 preprints describing new MinIon-based methods and results to preprint servers. For such a rapidly developing technology, the ability to share and consume methods from other scientists has created a feed forward effect where the technology has likely advanced at a faster rate than it otherwise would have.

Second, preprints have proven useful for rapidly correcting the scientific literature. On February 9, 2015, *Cell Systems* published a study that collected and analyzed metagenomic sequence data from the New York City subway system and reported finding *Yersinia pestis* and *Bacillus anthracis* (27). Because of the focus on these two bioterrorism agents, this study generated a considerable amount of media attention. On April 25, 2015, Petit et al. (28) posted a preprint to Zenodo demonstrating that there was no evidence for *B. anthracis* in the dataset. On July 29, 2015, a critique was published by *Cell Systems* along with a response from the original authors offering a correction to their manuscript (29, 30). A second anecdote of using preprints to aid in post-publication peer-review surrounds the publishing of a draft tardigrade genome in *The Proceedings of the National Academy of Sciences*. On November 23, 2015 a study by Boothby et al. (31) was published online. The authors claimed that 17.5% of its genes came from bacteria, archaea, fungi, plants, and viruses. Another group had been analyzing sequence data from a parallel tardigrade genome sequencing project and did not observe the same result. A week later, on December 1, 2015, the second group posted a preprint comparing the two genome sequences and demonstrating that the exciting claims of horizontal gene transfer were really the product of contaminants (32); this analysis would eventually be peer-reviewed and published online by the original journal on March 24, 2016 followed by a rebuttal by the original authors on May 31, 2016 (33, 34). Two other analyses of the original data were peer-reviewed and published in May 2016 and a third was posted as a preprint on February 2, 2016 (35–37). Both of these anecdotes underscore the value of having a rapid posting cycle to correcting errors in the scientific literature and that results posted to preprint servers were able to correct the record within weeks of the initial publication while the traditional peer-review path took six months in both cases. A final notable case where preprints have accelerated the correction of the scientific record was a preprint posted by Bik et al. (38) reporting numerous cases of image manipulation in peer-reviewed studies. This was a case where a journal may have been reluctant to publish the findings because it could have put the journal in a bad light. Posting the manuscript as a preprint removed potential conflicts of interests from journals that could have hindered its ability to be formally published in a journal. After the preprint was posted on April 20, 2016 it was peer-reviewed and published in *mBio* on June 7, 2016 (39). Instead of using preprints to react to published papers that have been through peer-review, it would be interesting to consider how the editorial process for these examples and the infamous “Arsenic Life” paper (40) would have been different had they initially been posted as preprints.

## Metrics for microbiology-affiliated preprints

To analyze the use of preprints, I downloaded the *bioRxiv* on April 17, 2017. I chose to analyze *bioRxiv* preprints because these preprints are amenable for submission to ASM journals and there were 9,780 *bioRxiv* preprints compared to the 2,911 preprints that were available at *PeerJ Preprint* on the same date. The code used to analyze these preprints and the rest of this manuscript are available as a reproducible GitHub repository at http://www.github.com/SchlossLab/Schloss_PrePrints_mBio_2017. Among the 9,780 preprints on bioRxiv, 483 were assigned by the authors into the Microbiology category. One limitation of the *bioRxiv* interface is the inability to assign manuscripts to multiple categories or to tag the content of the preprint. For example, this manuscript could be assigned to either the Microbiology or the Scientific Communication and Education categories. To counter this limitation, I developed a more permissive approach that classified preprints as being microbiology-affiliated if their title or abstract had words containing *yeast*, *fung*, *viral*, *virus*, *archaea*, *bacteri*, *microb*, *microorganism*, *pathogen*, or *protist*. I identified 1,617 additional manuscripts that I considered microbiology-affiliated. These microbiology-affiliated preprints were primarily assigned to the Evolutionary Biology (N=283), Bioinformatics (N=237), or Genomics (N=231) categories.

As the total number of preprints has grown exponentially since the creation of *bioRxiv*, submission of microbiology-affiliated preprints has largely followed this growth (Figure 1A). Although preprints are still relatively new, the collection of microbiology-affiliated preprints indicates widespread experimentation with the format and considerable geographic diversity. Reflecting the relative novelty of preprints, 1,484 (85.5%) corresponding authors who submitted a microbiology-affiliated preprint (N=1,735 total) have posted a single preprint and 4.6% have posted 3 or more preprints. Corresponding authors that have posted microbiology-affiliated preprints are from 67 countries and are primarily affiliated with institutions in the United States (46.2% of microbiology-affiliated preprints), United Kingdom (12.9%), and Germany (4.6%). As the preprint format matures, it will be interesting to see whether the fraction of authors that post multiple preprints increases and whether the geographic diversity amongst those authors is maintained.

**Figure 1.**
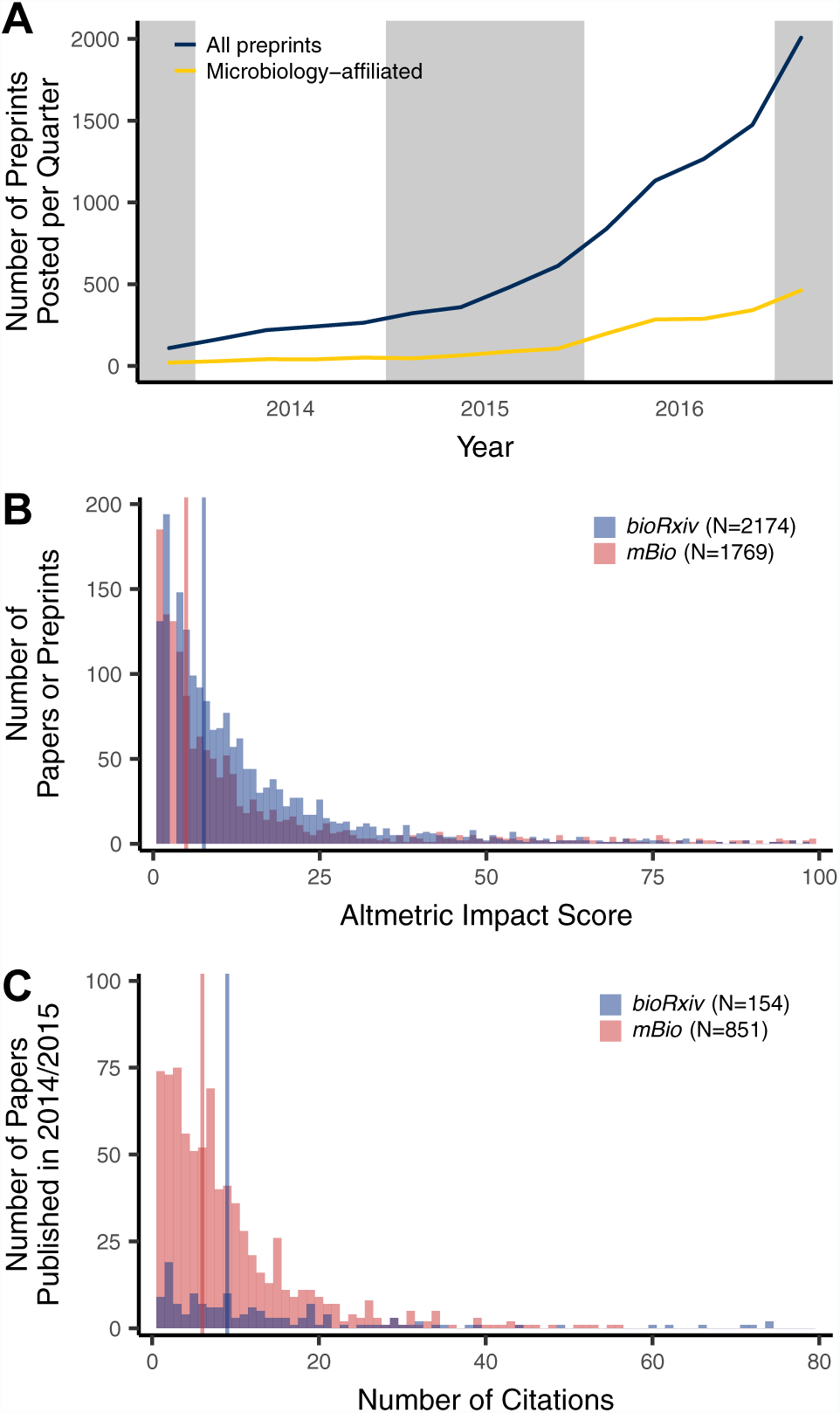
Summary of microbiology-affiliated preprints since the creation of *bioRxiv.* The total number of preprints posted for each quarter ending March 31, 2017 has largely tracked the overall submission of preprints to *bioRxiv* (A). The Altmetric attention scores of preprints posted to *bioRxiv* are similar to those published in *mBio* since November 2013 indicating preprints engender a similar level of attention (B). The number of times preprints that were published in 2014 and 2015 have been cited is similar to the number of citations for papers published in *mBio* in 2014 and 2015 indicates that published preprints are frequently cited (C). Regions with common background shading in A are from the same year. The vertical lines in B and C indicate the median Altmetric impact score and the median number of citations.

As stated above, preprints offer researchers the opportunity to improve the quality of their work by adding a more formal and public step to the scientific process. Among the microbiology-affiliated preprints, 197 (9.3%) had been commented on at least once and only 48 (2.3%) more than three times using the *bioRxiv*-hosted commenting feature. Although the hosted commenting is only one mechanism for peer-review, this result was somewhat disturbing since the preprint model implicitly depends on people’s willingness to offer others feedback. In spite of the lack of tradition within the scientific community to comment publicly online about colleagues’ research results, I am optimistic that this will change given the possibilities of new media (e.g. Twitter, Facebook, blogs); the advantage of the centralized commenting is that it is easier for the authors and others to integrate the feedback with the preprint. It is possible that incentives for open commenting and reviewing could shift the trend. Importantly, authors do appear to be incorporating feedback from colleagues or editorial insights from journals as 545 (25.9%) microbiology-affiliated preprints were revised at least once. Among the preprints posted prior to January 1, 2016, 31.3% of the Microbiology category preprints, 35.6% of the microbiology-affiliated preprints, and 33.6% of all preprints have been published. As noted above, not all authors submit their preprints to journals. This would indicate that the “acceptance rates” are actually higher. Regardless, considering that these acceptance rates are higher than many peer-reviewed journals (e.g. approximately 20% at ASM Journals), these results dispel the critique that preprints represent overly preliminary research.

Measuring the impact and significance of scientific research is notoriously difficult. Using several metrics I sought to quantify the effect that broadly defined microbiology-affiliated preprints have had on the work of others. Using the download statistics associated with each preprint, I found that the median number of times an abstract or PDF had been accessed was 922 (IQR: 601 to 1446) and 301 (IQR: 155 to 549), respectively. These values represent two aspects of posting a preprint. First, they reflect the number of times people were able to access science before it was published. Second, they reflect the number of times people were able to access a version of a manuscript that is published behind a paywall. To obtain a measure of a preprint’s ability to garner attention and engage the general public, I obtained the Altmetric Attention Score for each preprint (Figure 1B). The Altmetric Attention Score measures the number of times a preprint or paper is mentioned in social media, traditional media, Wikipedia, policy documents, and other sources; it does not include the number of citations (41). A higher score indicates that a preprint received more attention. Microbiology-affiliated preprints have had a median Altmetric Attention Score of 7.6 (IQR: 3.2 to 16.6) and those of all preprints hosted at *bioRxiv* have had a median score of 7.7 (IQR: 3.1 to 16.2). For comparison, the median Altmetric Attention Score for articles published in *mBio* published since 2013 was 5.0 (IQR: 1.5 to 14.5). Of all scholarship tracked by Altmetric, the median Altmetric Attention Score for preprints posted at *bioRxiv* ranks at the 87 percentile (IQR: 75 to 94). A controversial, yet more traditional metric of impact has been the number of citations an article receives. I obtained the number of citations for the published versions of manuscripts that were initially posted as preprints. To allow for a comparison to traditional journals, I considered the citations for preprints published in 2014 and 2015 as aggregated by Web of Science (Figure 1C). Among the preprints that were published and could be found in the Web of Science database, the median number of citations was 9 (IQR: 3-19; mean: 17.1). For comparison, among the papers published in *mBio* in 2014 and 2015, the median number of citations was 6 (IQR: 3-11; mean: 8.5). Although it is impossible to quantify the quality or impact of research with individual metrics, it is clear that the science presented in preprints and the publications that result from them are accepted by the microbiology community at a level comparable to more traditionally presented research.

## Preprints from an author’s perspective

Posting research as a preprint gives an author great control over when their work is made public. Under the traditional peer-review model, an author may need to submit and revise their work multiple times to several journals over a long period before it is finally published. In contrast, an author can post the preprint at the start of the process for others to consume and comment on as it works its way through the peer-review process. A first example illustrates the utility of preprints for improving access to research and the quality of its reporting. In 2014, my research group posted a preprint to *PeerJ Preprints* describing a method of sequencing 16S rRNA gene sequences using the Pacific Biosciences sequencing platform (42). At the same time, we submitted the manuscript for review at *PeerJ*. While the manuscript was under review, we received feedback from an academic scientist and from scientists at Pacific Biosciences that the impact of the results could be enhanced by using a recently released version of the sequencing chemistry. Instead of ignoring this feedback and resubmitting the manuscript to address the reviews, we generated new data and submitted an updated preprint a year later with a simultaneous submission to *PeerJ* that incorporated the original reviews as well as the feedback we received from the academic scientist and Pacific Biosciences. It was eventually published by *PeerJ* (43, 44). Since 2015, we have continued to post manuscripts as preprints at the same time as we have submitted manuscripts. Although the feedback to other manuscripts has not always been as helpful as our initial experience, in each case we were able to publicize our results prior to lengthy peer-review processes by immediately making our results available; in one case our preprint was available 7 months ahead of the final published version (45, 46). As another example, I posted a preprint of the current manuscript to *bioRxiv* on February 22, 2017. I then solicited feedback on the manuscript using social media. On March 14, 2017 I incorporated the comments and posted a revised preprint and submitted the manuscript to *mBio*. During that time, the abstract was accessed 189 times and the PDF was accessed 107 times. This process engaged 3 commenters on *bioRxiv*, 61 people either tweeted or re-tweeted the preprint on Twitter, 2 people on the manuscript’s GitHub repository, 1 person on a blog, and 2 via email. Compared to the two scientists that eventually reviewed the manuscript, the preliminary round of informal peer-review engaged a much larger and more diverse community than had I foregone the posting of a preprint. By the time that the final version of the manuscript was submitted on April 21, 2017, the preprint version of this manuscript had an Attention Score of 58, which placed it in the top 5% of all research scored by Altmetric and the abstract and PDF had been accessed 2,152 and 512 times, respectively. Although there are concerns regarding the quality of the science posted to a preprint server, I contend that responsible use of preprints as a part of the scientific process can significantly enhance the science.

## Preprints from a publisher’s perspective

A lingering question is what role traditional journals will have in disseminating research if there is broad adoption of preprints. Edited peer-reviewed journals offer and will continue to offer significant added value to a publication. A scholarly publishing ecosystem in which preprints coexist with journals will allow authors to gain value from the immediate communication of their work associated with preprints and also benefit from the peer-reviewed, professionally edited publication that publishers can provide.

The professional copyediting, layout, and publicity that these publishers offer are also unique features of traditional journals. An alternative perspective is that preprints will eventually replace traditional journals. Certainly, this is a radical perspective, but it does serve to motivate publishers to capture the innovation opportunities offered by preprints. By adopting preprint-friendly policies, journals can create an attractive environment for authors. As discussed above, a growing number of journals have created mechanisms for authors to directly submit preprints to their journals. An example is offered by the ASM, which earlier this year launched a new venture from *mSphere*. mSphereDirect is a publication track of the journal that capitalizes on the opportunity offered to couple preprints with rigorous peer-review. mSphereDirect actively encourages authors to post their manuscripts as preprints as part of an author-driven editorial process where an editorial decision is rendered within five days and publication in *mSphere* within a month (47). As the mSphereDirect mechanism evolves and is perhaps adopted by other journals, it will be interesting to see whether public feedback on preprints will be used to further streamline the editorial process. ASM is developing a new platform, MicroNow, which will help coalesce specific communities within the microbial sciences, further enhancing the use of preprints as well as published articles (Stefano Bertuzzi, personal communication). In addition to integrating preprints into the traditional editorial process, several professional societies have also explicitly supported citation of preprints in their other publications and recognize the priority of preprints in the literature (21–23). These are policies that empower authors and make specific journals more attractive. Other practices have great potential to improve the reputation of journals. As measured above, preprints are able to garner attention on par with papers published in highly selective microbiology journals. Thus, it is in a journal’s best interest to recruit these preprints to their journals. Several journals including *PLOS Genetics* and *Genome Biology* have publicly stated that they scout preprints for this purpose (48, 49). Preprints can also be viewed as a lost opportunity to journals. A preprint that garners significant attention may be ignored when it is finally published, bringing little additional attention to the journal. Going forward, there will likely be many innovative approaches that publishers develop to benefit from incorporating preprints into their process and whether publishers’ influence is reduced by the widespread adoption of preprints.

## Conclusions

Since the first microbiology-affiliated preprint was posted on bioRxiv in November 2013 (50), an increasing number of microbiologists are posting their unpublished work to preprint severs as an efficient method for disseminating their research prior to peer-review. A number of critical concerns remain about how widespread their adoption will be, how they will be perceived by traditional journals and other scientists, and whether traditional peer-review will adapt to the new scientific trends and technologies. Regardless, preprints should offer a great opportunity for both scientists and journals to publish high quality science.

## Acknowledgements

I am grateful to Stefano Bertuzzi and Lynn Enquist for their helpful comments on earlier versions of this manuscript and to the numerous individuals who provided feedback on the preprint version of the manuscript. This work was supported in part by funding from the National Institutes of Health (P30DK034933). I appreciate the support of Altmetric, Inc and Thompson Reuters who provided advanced programming interface (API) access to their databases. The workflow utilized commands in GNU make (v.3.81), GNU bash (v.4.1.2), and R (v.3.3.3). Within R I utilized the cowplot (v.0.7.0), dplyr (v.0.5.0), ggplot2 (v.2.2.1), httr (v.1.2.1), RCurl (v.1.95-4.8), rentrez (v.1.0.4), RJSONIO (v.1.3-0), rvest (v.0.3.2), sportcolors (v.0.0.1), and tidyr (v.0.6.1) packages. All journal policies and the information cited using webpage links was current on April 21, 2017.

---

**Supplemental Figure 1. Screen shot of the preprint for this manuscript at bioRxiv.**

**Supplemental Figure 2. Screen shot of a preprint by the author hosted at PeerJ Preprints.**

